# Bi-level diversity optimisation for representative protein panel selection

**DOI:** 10.64898/2026.04.17.719243

**Authors:** Zhen Ou, Katherine James, Simon Charnock, Anil Wipat

## Abstract

Selecting representative subsets from large protein sequence datasets is a common challenge in enzyme discovery and related tasks under limited screening capacity. In practice, candidate panels are often constructed using clustering-based redundancy reduction or manual selection guided by phylogenetic or similarity-network analyses, which do not directly optimise subset diversity and require threshold tuning or expert interpretation. Here, we present a bi-level diversity-optimisation framework for representative protein panel selection implemented using a local search heuristic that iteratively updates panel composition to improve diversity. The method formulates panel design as a combinatorial optimisation problem over pairwise distance matrices, combining a MaxMin objective to enforce minimum separation between selected sequences with a MaxSum objective to increase global dispersion. This formulation enables the direct construction of fixed-cardinality panels while remaining independent of the similarity representation used to compute pairwise distances. Benchmarking across four Pfam families shows that the bi-level formulation consistently reduces redundancy among selected sequences, lowering maximum pairwise identity by 43-46% relative to the previous MaxSum-based formulation, while maintaining comparable or improved EC-label coverage. The framework can incorporate sequence- or structure-based similarity measures, providing a flexible strategy for constructing diverse representative panels across homologous protein families.

## Introduction

Advances in high-throughput sequencing and metagenomics have greatly expanded enzyme sequence availability, enabling new opportunities for biocatalyst discovery and engineering. However, experimental characterisation remains constrained by fixed screening capacities, typically in plate-based formats. A central computational challenge is therefore to select a representative subset from a large homologous protein family that captures underlying protein diversity while avoiding redundancy.

Representative selection is most commonly addressed using clustering and redundancy-reduction tools such as CD-HIT [8], UCLUST [4], and MMSeqs2 [22]. These methods group sequences using predefined similarity thresholds and select a representative sequence from each cluster. While efficient, such approaches offer limited control over subset size and may retain near-duplicate sequences just below the chosen cut-off [9]. Phylogenetic and similarity network-based methods face similar challenges, including dependence on similarity thresholds, sensitivity to topology accuracy, and the need for manual interpretation [2, 6, 18, 14].

More generally, most existing approaches treat diversity as a by-product of clustering and are restricted to a single similarity definition, typically sequence identity. However, protein diversity can be defined across multiple feature spaces, including sequence similarity, structural similarity, or distances in learned representations. A general framework that directly optimises representative selection while remaining independent of the similarity representation is therefore desirable.

Optimisation-based strategies have recently been explored for representative selection. For example, submodular optimisation frameworks aim to maximise dataset coverage by selecting representatives such that each element remains close to at least one selected sequence [12]. While these approaches encourage diversity through coverage, they do not explicitly enforce separation among selected elements. Diversity problems provide a different class of optimisation models that formulate representative selection as a combinatorial optimisation task over pairwise distances [5]. In our previous work, we applied the MaxSum Diversity Problem (MSD) to enzyme panel design and showed that maximising total pairwise distance can recover diverse functional annotations when using global sequence identity as the distance metric [1]. However, because MSD maximises aggregate pairwise dispersion, it may still select closely related elements that are jointly distant from the remainder of the dataset.

Here we adopt a bi-level diversity formulation that combines the MaxMin (MMD) and MaxSum objectives as sequential optimisation levels [13]. In this context, the term ‘bi-level’ refers to a hierarchical combination of two objectives, rather than a nested leader–follower formulation used in classical bi-level optimisation. The MaxMin component enforces a minimum separation among selected elements, while the MaxSum component promotes global dispersion within that constraint. Using four Pfam families as benchmarks, we show that this formulation provides a practical framework for selecting diverse and stable protein panels that capture broad functional annotations across multiple similarity representations.

## Methods

### Problem Formulation

The optimisation framework builds on the DropAdd Tabu Search algorithm (DropAdd-TS) of Porumbel et al. (2011)[15]. Let *D* = (*d*_*ij*_ ) denote a pairwise distance matrix derived from similarity scores *s*_*ij*_, where *d*_*ij*_ = 1 − *s*_*ij*_ . Given a candidate set *V* and a target subset size *k*, the MMD diversity objective seeks a subset *S* ⊂ *V*, |*S*| = *k*, that maximises the minimum pairwise distance:

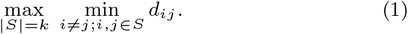

To increase global dispersion among already separated elements, a secondary MSD objective maximises the total pairwise distance:

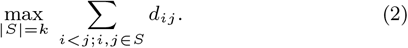

The objectives are evaluated hierarchically: the MMD objective defines the primary optimisation criterion, and the MSD objective acts as a secondary criterion among solutions with equal minimum distance.

### Optimisation Strategy

An initial solution is constructed greedily. The first element is selected as the vertex with the largest total distance to all other vertices. Subsequent elements are added by maximising the minimum distance to the current solution (MMD objective). When ties occur, the candidate with the largest summed distance to the current solution is selected (MSD tie-breaking).

The main DropAdd-TS search iteratively removes the oldest element in *S* and inserts a new candidate from *V \ S* according to the same hierarchical MMD-MSD criterion. For each vertex *i* ∈ *V*, the minimum distance to *S* (MinDist(*i, S*)), and the total distances to *S* (SumDist(*i, S*)) are maintained and updated to rank candidates. To reduce neighbourhood size, candidate selection is restricted to the top 50 add moves ranked by MinDist.

To improve robustness and computational efficiency, a dynamic move-based tabu list was adopted, with lower and upper bounds proportional to subset and superset size. The tabu tenure increases during stagnation and resets upon improvement. If stagnation persists at maximum tenure, a random add move is performed.

A solution is considered improved if it increases the MMD objective, or if the MMD objective is unchanged and the MSD objective increases. Tabu moves are allowed if the resulting subset improves the current best solution *S*^*^, and the oldest tabu move is permitted when no admissible moves remain to ensure that a valid add move is always performed. The search terminates after 2,000 consecutive non-improving iterations (scaled with problem size up to 20,000), or after 50,000 steps (10*n* for large instances) (Algorithm 1).

#### Algorithm 1

Bi-level DropAdd Tabu Search

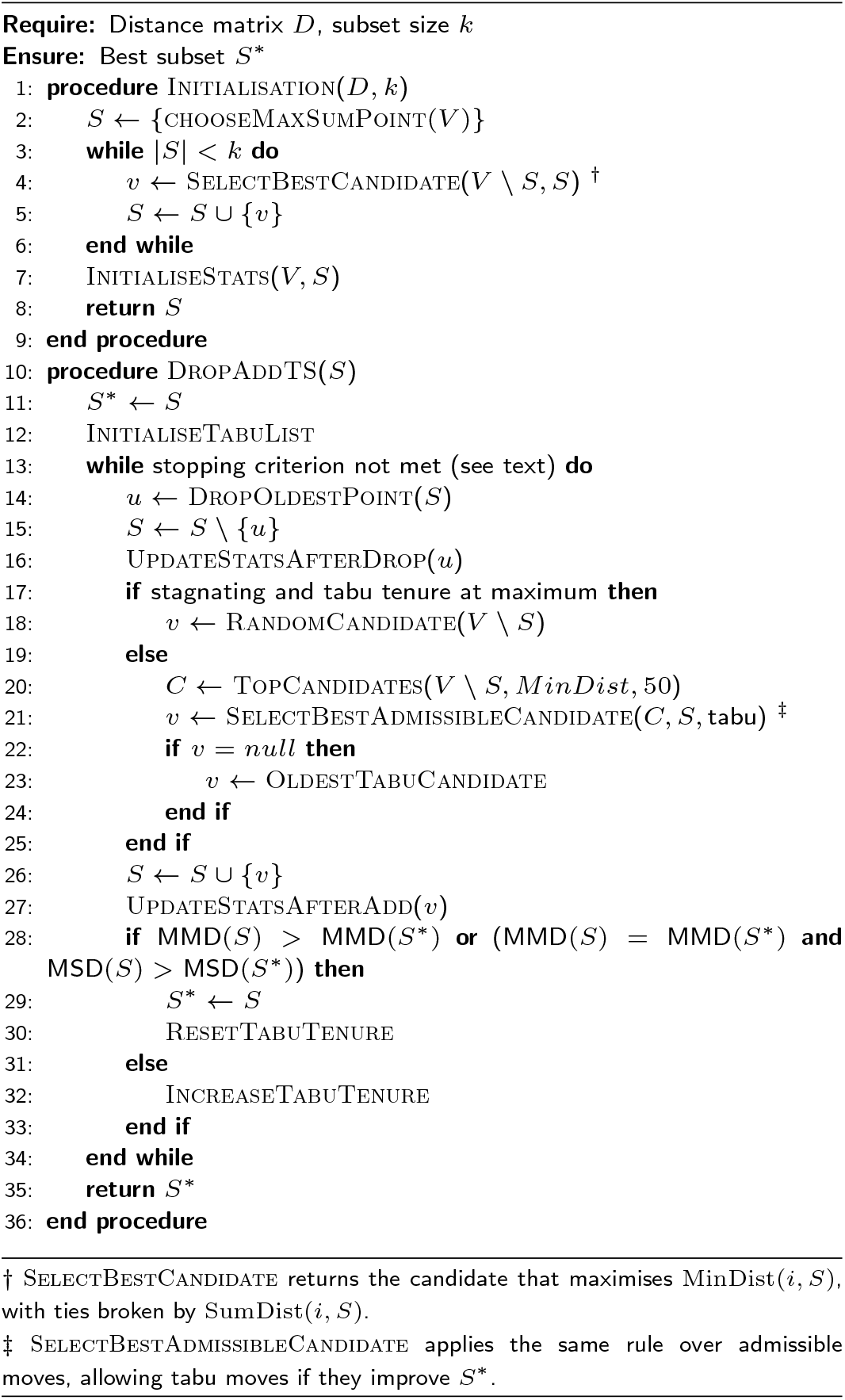

### Dataset Collection

Protein sequences were obtained from the Swiss-Prot section of UniProt (2024 03 release; [23]) and filtered according to three criteria: (i) bacterial origin, (ii) presence of a Pfam classification, and (iii) complete Enzyme Commission (EC) annotation.

Sequences were grouped by Pfam family to generate homologous datasets. Families containing more than 900 sequences and at least 20 unique EC labels were retained. These thresholds were chosen to ensure sufficient size and functional diversity for meaningful comparison between selection strategies.

To minimise the effect of length variation on pairwise similarity scores, sequences deviating from the family mean length were removed using family-specific standard deviations (Table 7). Families falling below the sequence or EC thresholds after length filtering were excluded.

Predicted structures for the retained sequences were retrieved from AlphaFoldDB [25], and sequences without available models were removed to ensure consistency between sequence- and structure-based analyses.

Sequences annotated with multiple EC numbers were treated as distinct functional categories in this work, with each unique EC label combination considered a separate class (Table 1). The selected families include both enzyme and non-enzyme proteins, with the ABC transporter family illustrating the broader applicability of the method beyond enzyme-specific contexts.

**Table 1.** Summary statistics of the four Pfam subsets after filtering. Columns report the Pfam accession, domain name, number of sequences, number of EC labels, and the original Gini-Simpson index (GSI) of the dataset.

| Pfam ID | Pfam name | Sequences | EC | GSI |
| --- | --- | --- | --- | --- |
| PF04055 | Radical SAM superfamily (SAM) | 2872 | 26 | 0.835 |
| PF00005 | ABC transporter (ABC) | 1442 | 31 | 0.891 |
| PF00171 | Aldehyde dehydrogenase family (ADH) | 959 | 43 | 0.679 |
| PF00155 | Aminotransferase class I and II (ATF) | 934 | 28 | 0.652 |

In addition to the curated Pfam datasets used for benchmarking, a highly redundant metagenomic aminotransferase dataset (996 sequences) provided by Prozomix Ltd. was used to illustrate representative selection behaviour in dense sequence similarity networks. This dataset was used solely for visualisation purposes and was not included in the quantitative benchmarking analyses.

### Diversity and Stability Metrics

Functional diversity of selected subsets was evaluated using two complementary metrics: Label Coverage (LC) and the Gini-Simpson Index (GSI).

LC quantifies the proportion of unique EC labels present in a selected subset relative to the total number of EC labels in the original dataset, ranging from 0 (no labels captured) to 1 (complete coverage). The GSI quantifies diversity by incorporating both label richness and evenness, representing the probability that two randomly selected sequences belong to different labels [3]. Higher GSI values indicate more balanced distributions of functional classes, with a maximum value of 1 when labels are uniformly represented.

To assess selection stability across stochastic runs, pairwise Jaccard indices were computed at both the sequence and EC-label levels. The index measures the overlap between subsets as the ratio of their intersection to their union, with values ranging from 0 (no overlap) to 1 (identical subsets).

### Feature Representations

#### Sequence identity and similarity

Pairwise global alignments were computed using EMBOSS Needleall (v6.6.0.0; [7, 16]) with the BLOSUM62 substitution matrix. Needleall [11] reports both sequence identity (fraction of exact residue matches) and sequence similarity (fraction of identical or conservatively substituted positions), normalised by aligned length to reflect full-length similarity.

Pairwise local identities were additionally computed using Smith-Waterman alignments (EMBOSS water; [21]), which capture conserved regions shared between sequences, and normalised by the longer sequence length to avoid inflation from short high-scoring regions. Local alignment-based identities were included to support comparison with clustering-based methods that rely on local sequence similarity.

#### Structural similarity

Pairwise structural similarity was computed using TM-align (v20170708; [27]). For each protein pair, the final similarity was defined as the average of the two directional TM-scores (each normalised by the length of one protein), providing a symmetric global fold similarity measure.

### Computational Tools and Baselines

TS-MA [1] was used as the reference diversity-optimisation algorithm for the MSD objective with parameters as originally reported. To provide a baseline for the MMD objective, a greedy MaxMin heuristic was also evaluated. This method follows the same construction procedure used to initialise DropAdd-TS, iteratively adding the candidate that maximises the minimum distance to the current subset. To introduce stochasticity comparable to the other optimisation methods, the initial element was selected randomly.

Performance was evaluated over 50 independent runs for each subset size and dataset. For experiments involving subset expansion, DropAdd-TS was similarly executed over 50 independent runs, initialised from the best solution obtained at the smaller subset sizes (*k* = |*L*| or *k* = 2|*L*|), where |*L*| denotes the number of EC labels in the dataset.

CD-HIT (v4.8.1; [8]) and MMseqs2 (v15.6f452; [22]) were used for sequence-based clustering, with identity thresholds adjusted to obtain subsets of comparable sizes. Foldseek (v9.427df8a; [24]) was used for structure-based clustering with TM-score thresholds tuned similarly. Identity and TM-score thresholds used are reported in Table 10.

DropAdd-TS was implemented in Python 3.11 within a Nextflow workflow. Experiments were conducted on a 1.4 GHz Intel Core i5 CPU with 8GB RAM. Similarity networks were visualised using Cytoscape (v3.10.3; [20]) with an organic layout, where nodes represent proteins and edges connect sequence pairs exceeding the specified similarity threshold.

## Results

### Diversity-based Algorithm Comparison

We compared functional diversity, redundancy control, and selection stability across four Pfam families for subsets of size *k* = 100 (Table 2-3). Differences between methods were not formally tested for statistical significance and should be interpreted in the context of variability across runs.

**Table 2.** Functional diversity and redundancy control for subsets of size *k* = 100 across four Pfam families. LC: label coverage; GSI: Gini-Simpson index; Max_ID: maximum pairwise sequence identity within each selected subset. Results are reported as mean ± standard deviation over 50 independent runs. Best values per metric within each Pfam family are shown in bold (higher is better for LC and GSI, lower is better for Max_ID).

| Pfam | Method | LC | GSI | Max.ID |
| --- | --- | --- | --- | --- |
| SAM | Random | 0.345±0.055 | 0.826±0.012 | 0.999±0.002 |
| | Greedy | 0.962±0.000 | 0.89±0.002 | 0.37±0.003 |
| | TS-MA | **0.993±0.015** | **0.917±0.002** | 0.661±0.008 |
| | DropAdd-TS | 0.962±0.000 | 0.891±0.001 | **0.355±0.001** |
| ABC | Random | 0.576±0.05 | 0.883±0.015 | 1.0±0.001 |
| | Greedy | 0.93±0.028 | 0.935±0.003 | 0.412±0.003 |
| | TS-MA | 0.886±0.032 | **0.941±0.002** | 0.734±0.045 |
| | DropAdd-TS | **0.951±0.019** | 0.938±0.001 | **0.4±0.001** |
| ADH | Random | 0.304±0.046 | 0.666±0.044 | 1.0±0.001 |
| | Greedy | 0.86±0.018 | 0.816±0.006 | 0.522±0.003 |
| | TS-MA | 0.795±0.024 | **0.855±0.005** | 0.894±0.000 |
| | DropAdd-TS | **0.901±0.011** | 0.823±0.001 | **0.509±0.000** |
| ATF | Random | 0.356±0.067 | 0.648±0.026 | 0.999±0.002 |
| | Greedy | 0.764±0.04 | 0.723±0.01 | 0.417±0.003 |
| | TS-MA | 0.734±0.029 | **0.84±0.004** | 0.762±0.006 |
| | DropAdd-TS | **0.774±0.025** | 0.734±0.001 | **0.408±0.000** |

Random selection performed poorly across all datasets, capturing only 30-58% of EC labels, exhibiting near-zero sequence-level Jaccard overlap across runs, and producing maximal pairwise identities approaching 1.0.

TS-MA substantially improved performance relative to random selection and achieved the highest GSI values across all datasets, indicating more balanced label distributions. However, because the method optimises only a MSD objective, redundancy was only partially controlled. In three of the four families, TS-MA subsets still contained highly identical sequence pairs (Max ID ≥ 0.73).

The baseline greedy MMD method and DropAdd-TS produced comparable LC values, underscoring the value of enforcing a minimum separation constraint. DropAdd-TS achieved the lowest maximum pairwise identity across all families, indicating stronger control over closely related elements than MMD alone. Notably, these results were obtained without prior redundancy reduction, showing that DropAdd-TS effectively limits internal similarity. Furthermore, DropAdd-TS obtained the highest LC in three of four families, missing only a single label in the remaining dataset.

The higher LC achieved by DropAdd-TS was accompanied by lower GSI values relative to TS-MA, reflecting a trade-off between label coverage and distribution evenness. As subset size increases, additional representatives are more likely drawn from dominant functional classes, increasing LC while reducing label balance. This effect is more pronounced in skewed datasets such as ADH and ATF, where dominant EC labels account for a large proportion of the sequences.

DropAdd-TS also exhibited the highest sequence-level Jaccard overlap across independent runs in all families, indicating consistent convergence toward similar subsets (Table 3). While TS-MA showed marginally higher EC-label stability in two datasets, one of these cases (ATF) coincided with lower LC, suggesting convergence toward a less diverse functional representation.

**Table 3.** Stability of subset selection across 50 independent runs for subsets of size *k* = 100. Stability is measured using the mean pairwise Jaccard index at the sequence level (Jaccard (Seq)) and EC-label level (Jaccard (EC)). Higher values indicate greater consistency across runs. Best values per metric within each Pfam family are shown in bold.

| Pfam | Method | Jaccard (Seq) | Jaccard (EC) |
| --- | --- | --- | --- |
| SAM | Random | 0.018±0.009 | 0.667±0.108 |
| | Greedy | 0.311±0.045 | 0.963±0.038 |
| | TS-MA | 0.444±0.05 | **0.988±0.018** |
| | DropAdd-TS | **0.581±0.075** | 0.977±0.035 |
| ABC | Random | 0.036±0.013 | 0.733±0.074 |
| | Greedy | 0.407±0.056 | 0.94±0.039 |
| | TS-MA | 0.522±0.045 | 0.93±0.038 |
|  | DropAdd-TS | **0.639±0.09** | **0.975±0.022** |
| ADH | Random | 0.056±0.016 | 0.472±0.081 |
| | Greedy | 0.565±0.057 | 0.954±0.033 |
| | TS-MA | 0.576±0.039 | 0.934±0.038 |
|  | DropAdd-TS | **0.793±0.077** | **0.98±0.026** |
| ATF | Random | 0.057±0.016 | 0.478±0.11 |
| | Greedy | 0.444±0.062 | 0.852±0.061 |
| | TS-MA | 0.537±0.045 | **0.921±0.042** |
| | DropAdd-TS | **0.711±0.071** | 0.908±0.055 |

Runtime increased with problem size for both optimisation methods. For the largest family (SAM), DropAdd-TS required longer runtimes (∼231s) than TS-MA (∼40s). For the smaller families, runtimes were comparable, with DropAdd-TS ranging from 19-46s and TS-MA from 16-31s (Table 8).

To complement these quantitative metrics, the selection behaviour of both algorithms was visualised using a sequence similarity network. Subsets of size *k* = 100 were projected onto a network constructed from a highly redundant metagenomic aminotransferase dataset (996 sequences) at an 80% identity threshold, chosen to group highly similar sequences while preserving cluster structure (Fig. 1). While both algorithms sample representatives from multiple regions of the network, TS-MA occasionally includes multiple sequences from densely connected clusters. In contrast, DropAdd-TS distributes representatives more evenly across clusters, typically selecting a single sequence per group. This behaviour is consistent with the lower maximum pairwise identity observed in the DropAdd-TS subsets and reflects the minimum-distance constraint imposed by the MMD objective.

**Figure 1.**
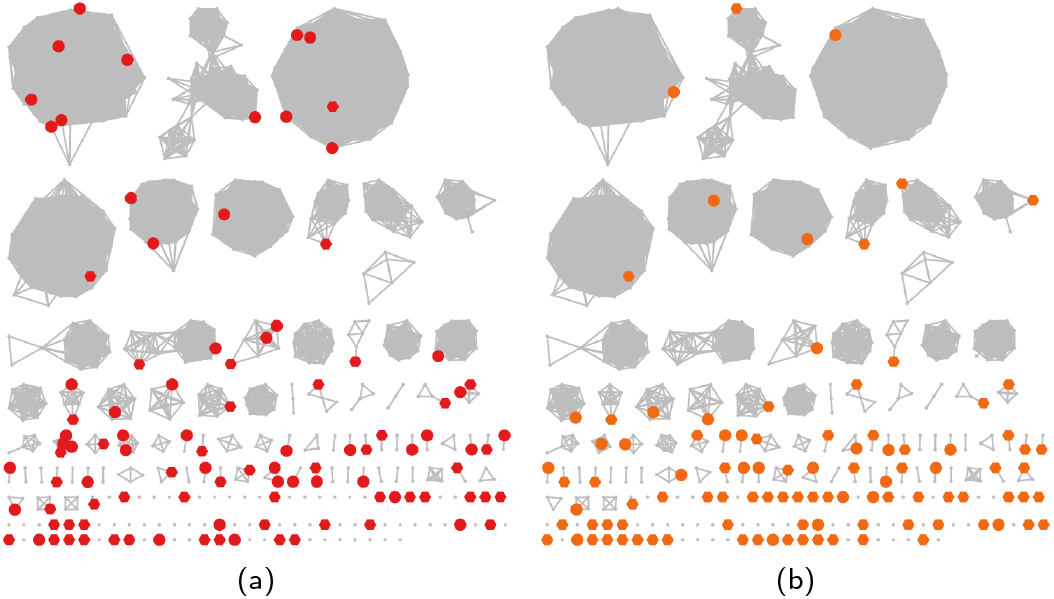
Representative selections projected onto a sequence similarity network (80% identity threshold; 996 sequences; *k* = 100). Grey nodes represent sequences, and highlighted nodes indicate representatives selected by (a) TS-MA and (b) DropAdd-TS.

Overall, DropAdd-TS provides an efficient alternative approach with improved redundancy control and convergence stability relative to single-level diversity optimisation, while maintaining competitive recovery of EC label annotations.

### Comparison with Clustering Methods

DropAdd-TS was next evaluated against the widely used sequence-based clustering tools CD-HIT [8] and MMseqs2 [22]. Because clustering methods do not directly control subset cardinality, identity thresholds were adjusted to produce representative sets close to *k* ≈ 100. DropAdd-TS was then executed over 50 runs for the same subset sizes obtained from MMSeqs2. Local alignment-based identities were additionally evaluated to enable direct comparison with the similarity measures used by these clustering methods.

DropAdd-TS produced subsets with average LC broadly comparable to those obtained by CD-HIT and MMSeqs2 (Table 4). Although LC was slightly lower in some datasets, it matched or exceeded the clustering methods in the others and was accompanied by considerably lower maximum pairwise sequence identity. This pattern is illustrated in Fig. 2, where the best achievable DropAdd-TS solutions generally occupy regions of lower sequence identity while maintaining similar or higher EC label coverage.

**Table 4.** Functional diversity and redundancy control of representative subsets from four Pfam families selected using CD-HIT, MMseqs2, and DropAdd-TS. LC: label coverage; GSI: Gini-Simpson index; Max_ID: maximum pairwise sequence identity within each subset. DropAdd-TS results are reported as mean ± standard deviation across 50 runs. Best values per metric are shown in bold (higher is better for LC and GSI, lower is better for Max_ID).

| Pfam | Method | LC | GSI | Max_ID |
| --- | --- | --- | --- | --- |
| SAM | CD-HIT | **0.962** | 0.886 | 0.465 |
|  | MMSeqs2 | **0.962** | 0.891 | 0.527 |
|  | DropAdd-TS (global) | **0.962±0.0** | **0.893±0.001** | **0.353±0.001** |
|  | DropAdd-TS (local) | **0.962±0.0** | **0.891±0.0** | **0.364±0.001** |
| ABC | CD-HIT | 0.935 | 0.932 | 0.442 |
|  | MMSeqs2 | **0.968** | **0.943** | 0.479 |
| | DropAdd-TS (global) | 0.955±0.017 | 0.937±0.001 | **0.401±0.0** |
| | DropAdd-TS (local) | 0.965±0.009 | 0.937±0.001 | 0.408±0.001 |
| ADH | CD-HIT | 0.86 | **0.834** | 0.528 |
|  | MMSeqs2 | 0.86 | 0.83 | 0.557 |
| | DropAdd-TS (global) | **0.901±0.011** | 0.823±0.001 | **0.509±0.0** |
| | DropAdd-TS (local) | 0.881±0.008 | 0.822±0.002 | 0.512±0.0 |
| ATF | CD-HIT | **0.821** | 0.735 | 0.45 |
|  | MMSeqs2 | **0.821** | 0.733 | 0.463 |
| | DropAdd-TS (global) | 0.774±0.025 | 0.734±0.001 | **0.408±0.0** |
| | DropAdd-TS (local) | 0.795±0.021 | **0.743±0.001** | 0.409±0.0 |

**Figure 2.**
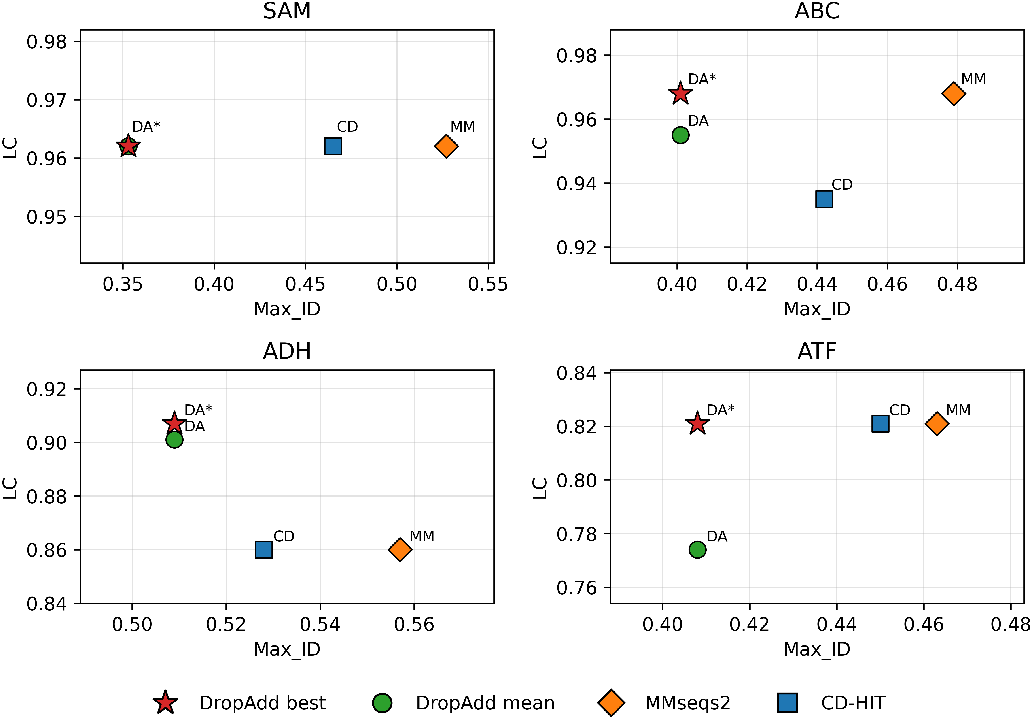
Label coverage (LC) and maximum pairwise sequence identity (Max ID) of representative subsets selected by CD-HIT (CD), MMseqs2 (MM), and DropAdd-TS (global sequence identity) across four Pfam families. Green circles denote the mean DropAdd-TS performance across 50 runs (DA), while red stars denote the best solution obtained (DA*).

Clustering-based methods reduce redundancy through fixed identity thresholds, below which representatives may still remain relatively close in sequence space. In contrast, the optimisation objective of DropAdd-TS prioritises representatives that are maximally separated, achieving similar LC while promoting broader coverage of sequence diversity.

Results obtained using local sequence identities were broadly consistent with those computed from global alignments. In the ABC and ATF families, LC was marginally higher when local alignment scores were used, while the relative performance of the methods remained unchanged. Local alignment may be preferable when proteins share conserved domains but differ in overall architecture, whereas global alignment is more appropriate when full-length similarity is representative of functional relationships.

To examine how functional classes were redistributed during representative selection, the per-label contributions to the GSI from the top 12 EC labels were compared between the selected subsets and the original datasets (Fig. 3). For DropAdd-TS, the best solution obtained using global sequence identity was used for comparison.

**Figure 3.**
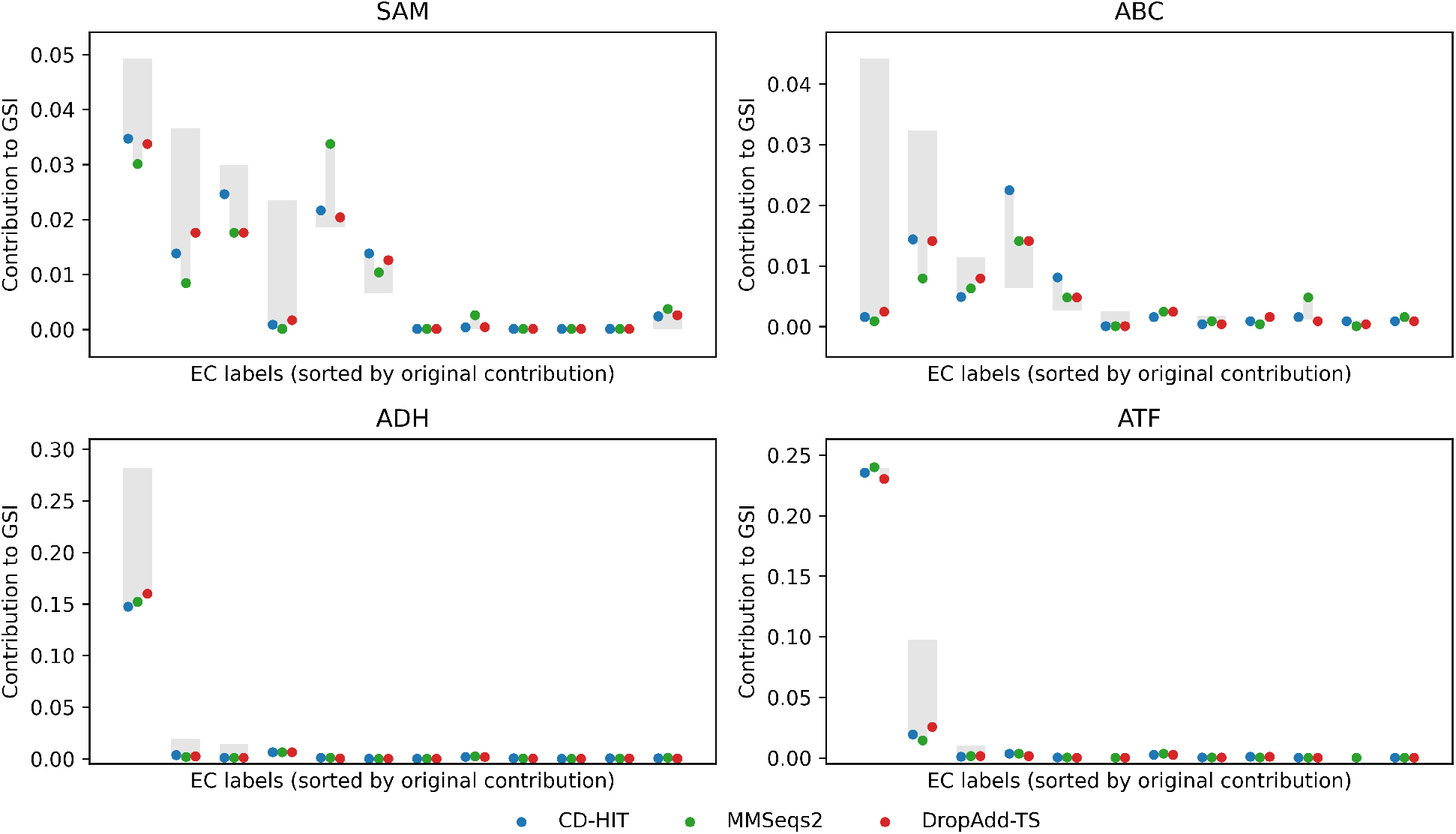
Redistribution of the top 12 EC label contributions to the Gini-Simpson index (GSI) for representative subsets of size ∼100 selected by CD-HIT, MMSeqs2, and DropAdd-TS across four Pfam families. Grey bars indicate the contribution of each EC label in the original dataset, with labels sorted by decreasing contribution. Coloured points show the corresponding contributions in the selected subsets.

In the two most skewed families (ADH and ATF), all three methods produced similar redistribution patterns, with dominant EC labels remaining strongly represented and resulting in lower GSI values (Table 4). In contrast, the more balanced families (SAM and ABC) exhibited greater variation between methods. Labels with higher within-label sequence identity tended to collapse into fewer representatives even when highly abundant in the dataset. Despite quantitative differences, all approaches generally adjusted the same labels in the same direction relative to the original distribution, indicating similar overall selection behaviour.

Although the overall redistribution trends were comparable, DropAdd-TS tended to produce slightly less extreme changes in label representation relative to clustering-based methods. Together with the lower maximum pairwise identity observed in Table 4, these results suggest that our diversity-optimised selection preserves EC label coverage and maintains balanced label distributions comparable to redundancy-based clustering, while further increasing separation among the selected sequences.

### Subset Expansion

To support incremental panel growth, an expansion mode was implemented in DropAdd-TS that selects additional representatives while accounting for previously selected members.

Across all Pfam families, subsets obtained through expansion achieved higher LC than subsets generated directly at the same target size (Fig. 4). However, this improvement was accompanied by a reduction in evenness and an increase in maximum pairwise identity, reflecting the reduced search space once part of the sequence diversity has already been selected.

**Figure 4.**
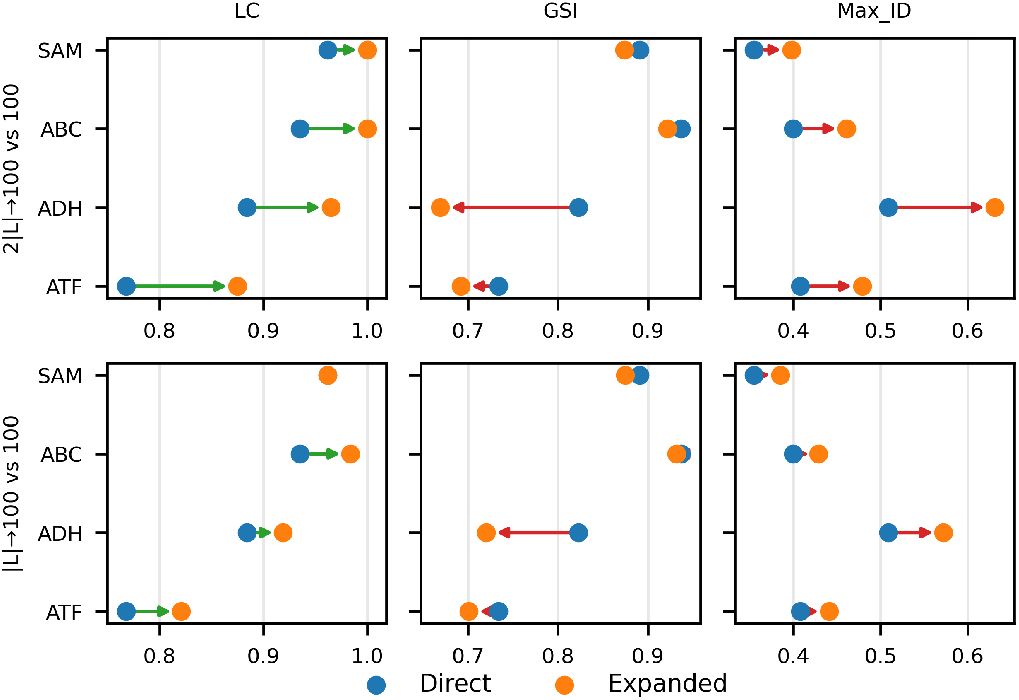
Impact of incremental subset expansion on functional diversity metrics across four Pfam families. Blue points represent subsets generated directly at the target size and orange points represent subsets obtained by expansion; arrows indicate the change between direct selection and expansion. Expansion results are shown as mean values across 50 independent runs. Expansion generally increases label coverage (LC) but tends to reduce evenness (GSI) and increases maximum pairwise sequence identity (Max_ID).

The expansion interval also influenced the outcome. Direct expansion from *k* = |*L*| to *k* = 100 preserved greater flexibility to explore the remaining sequence diversity, resulting in higher evenness and lower redundancy compared to the more restrictive 2|*L*| → 100 expansion. In contrast, the constrained 2|*L*| → 100 expansion consistently achieved higher LC, reaching complete coverage for SAM and ABC. This effect arises because sequences selected in earlier iterations limit the remaining search space, favouring candidates that are more distant from the current subset. When sequence distance coincides with EC-labelled differences, previously unrepresented labels are more likely to be recovered.

### Multi-Feature Evaluation

Diversity-based models operate on generalized distance matrices and are therefore not constrained to sequence identity alone. To evaluate this flexibility, DropAdd-TS was applied using alternative feature representations, including sequence similarity derived from BLOSUM62 substitution scores and structural similarity based on TM-scores, in addition to the default global sequence identity.

Across all Pfam families, identity-based representations consistently achieved the highest LC and GSI values, suggesting that sequence identity provides the most effective separation of proteins when functional diversity is approximated using EC annotations (Table 5). Identity-based optimisation also produced the lowest maximum pairwise similarity, reflecting the greater separation between sequences in identity-based representations.

**Table 5.**
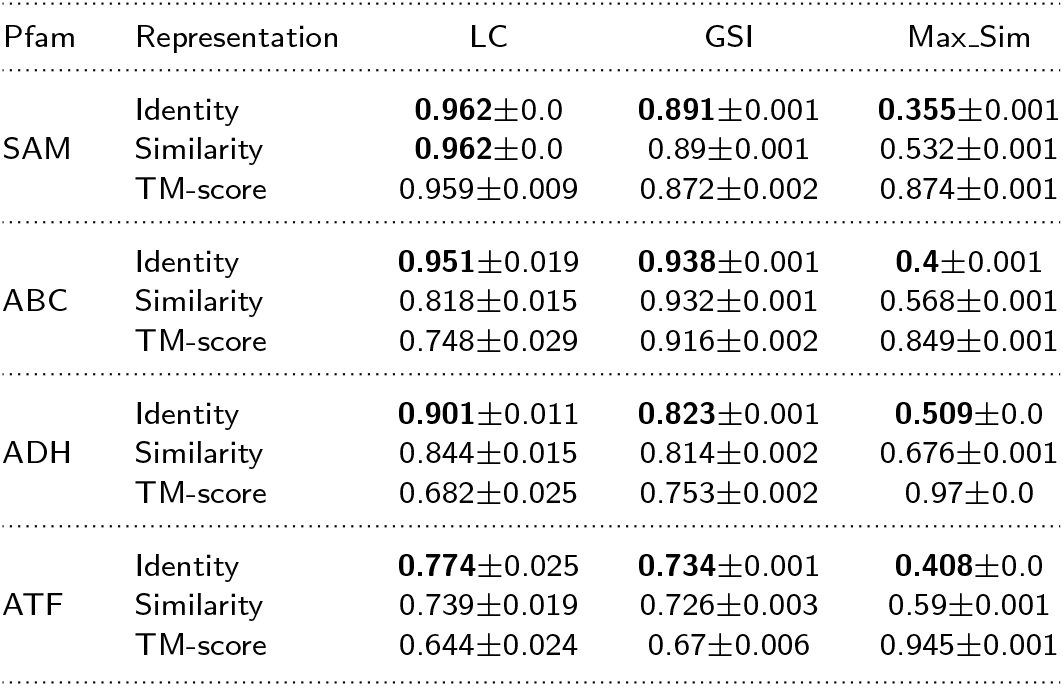
Performance of DropAdd-TS across different feature representations: global sequence identity, BLOSUM62-based sequence similarity, and global structural similarity (TM-score). Metrics reported are LC (label coverage), GSI (Gini-Simpson index), and Max_Sim (maximum pairwise similarity within each subset). Results are reported as mean ± standard deviation across 50 runs. Best values per dataset per metric are shown in bold (higher is better for LC and GSI, lower is better for Max_Sim).

Substituting identity with BLOSUM62-based similarity resulted in modest reductions in LC and GSI, along with an increase in Max Sim as expected. Since similarity scoring treats conservative substitutions as equivalent, sequences with relatively low identities may still appear close in similarity space, potentially reducing the separation between proteins associated with different EC labels. However, the effect was not uniform across datasets. In the SAM family, similarity-based representations achieved the same LC and only a marginally lower GSI compared with identity-based optimisation.

In contrast, structural similarity derived from TM-scores resulted in the largest decline in LC and GSI across all datasets. This behaviour likely reflects the strong evolutionary conservation of protein folds within Pfam families. While primary sequences can diverge substantially, their tertiary structures often remain similar to preserve stability and functions [26]. As a result, structural representations compress the available search space and limit the algorithm’s ability to separate representatives belonging to distinct functional annotations, which is also reflected in the substantially higher Max Sim values observed for TM-score-based optimisation.

To further examine this limitation, DropAdd-TS was compared with Foldseek [24], a fast structure-based clustering method. TM-score thresholds were adjusted in Foldseek to produce representative sets close to *k* ≈ 100. DropAdd-TS was then executed for the same subset sizes over 50 runs (Table 6).

**Table 6.** Comparison of representative subsets selected using DropAdd-TS and Foldseek under TM-score-based structural similarity. Metrics reported are LC (label coverage), GSI (Gini-Simpson index), and Max_TM (maximum pairwise TM-score within each subset). DropAdd-TS results are reported as mean ± standard deviation across 50 runs, with the best solution by label coverage shown in parentheses. Best values per metric are shown in bold (higher is better for LC and GSI, lower is better for Max_TM).

| Pfam | Method | LC | GSI | Max_TM |
| --- | --- | --- | --- | --- |
| SAM | Foldseek | **0.962** | **0.891** | 0.953 |
| | DropAdd-TS | 0.952±0.017 (0.962) | 0.869±0.003 | **0.873**±0.001 |
| ABC | Foldseek | **0.806** | 0.912 | 0.951 |
| | DropAdd-TS | 0.748±0.029 (0.806) | **0.916**±0.002 | **0.849**±0.001 |
| ADH | Foldseek | **0.744** | **0.804** | 0.986 |
| | DropAdd-TS | 0.675±0.023 (0.721) | 0.758±0.001 | **0.97**±0.0 |
| ATF | Foldseek | 0.643 | **0.715** | 0.975 |
| | DropAdd-TS | **0.666**±0.017 (0.679) | 0.672±0.007 | **0.946**±0.0 |

**Table 7.** Summary statistics of the four Pfam datasets before and after sequence-length filtering. The table reports dataset size, number of EC labels, sequence-length variation, the applied percentile-based length filters, and the resulting dataset sizes after length filtering only. Final dataset sizes used in benchmarking are reported in Table 1 after additional removal of sequences lacking AlphaFoldDB models.

| Pfam ID | Pfam name | Seqs | EC | Length SD | Filter* | Seq (filtered) | EC (filtered) |
| --- | --- | --- | --- | --- | --- | --- | --- |
| PF04055 | Radical SAM superfamily (SAM) | 3163 | 32 | 55 | 5%-95% | 2873 | 26 |
| PF00005 | ABC transporter (ABC) | 2181 | 44 | 63 | 17.5%-82.5% | 1443 | 31 |
| PF00171 | Aldehyde dehydrogenase family (ADH) | 1045 | 48 | 34 | 5%-95% | 959 | 43 |
| PF00155 | Aminotransferase class I and II (ATF) | 954 | 31 | 19.5 | 1%-99% | 935 | 28 |

\*Length ranges corresponding to these filters were: SAM (298-483 aa), ABC (253-503 aa), ADH (414-514 aa), and ATF (345-457 aa).

**Table 8.** Runtime comparison between TS-MA and DropAdd-TS across the four Pfam datasets. Values are reported as mean ±standard deviation across 50 runs.

| Pfam | Sequences | TS-MA (s) | DropAdd-TS (s) |
| --- | --- | --- | --- |
| SAM | 2872 | 40.983±8.72 | 231.383±87.601 |
| ABC | 1442 | 18.941±0.923 | 46.615±11.732 |
| ADH | 959 | 31.104±2.116 | 19.279±4.646 |
| ATF | 934 | 16.176±0.887 | 21.219±5.914 |

**Table 9.**
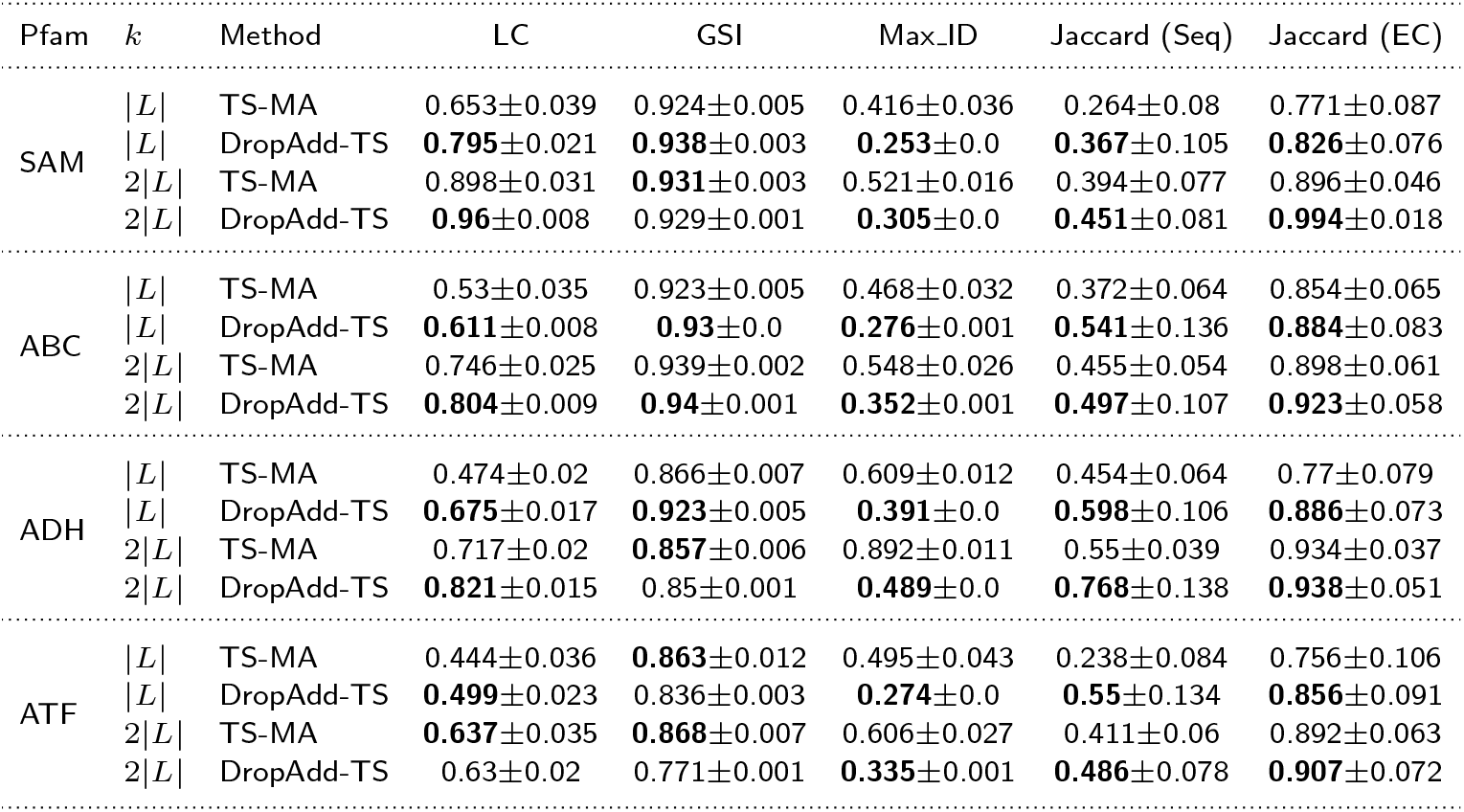
Comparison of TS-MA and DropAdd-TS for representative subsets across four Pfam families at two subset sizes (*k* = |*L*| and *k* = 2|*L*|). LC: label coverage; GSI: Gini-Simpson index; Max_ID: maximum pairwise sequence identity within each selected subset. Results are reported as mean ± standard deviation over 50 independent runs. Best values per metric within each Pfam family and subset size are shown in bold (higher is better for LC and GSI, lower is better for Max_ID).

**Table 10.** Clustering thresholds used to obtain representative subsets close to *k* ≈ 100 for CD-HIT, MMseqs2, and Foldseek across the four Pfam datasets. The resulting subset sizes (*k*) are reported for each method.

| Pfam | CD-HIT identity | k | MMSeqs2 identity | k | Foldseek TM-score | k |
| --- | --- | --- | --- | --- | --- | --- |
| SAM | 0.4 | 102 | 0.449 | 98 | 0.935 | 99 |
| ABC | 0.446 | 100 | 0.49 | 101 | 0.902 | 100 |
| ADH | 0.532 | 99 | 0.571 | 100 | 0.986 | 97 |
| ATF | 0.437 | 101 | 0.473 | 100 | 0.967 | 103 |

Consistent with the trends observed above, Foldseek produced lower LC and GSI values than sequence-based clustering methods. While Foldseek achieved higher LC and GSI than the average DropAdd-TS performance under structural similarity, the best achievable DropAdd-TS solutions reached LC values comparable to those obtained by Foldseek in most datasets. Again, DropAdd-TS consistently produced subsets with lower maximum pairwise TM-scores, imposing greater separation among the selected representatives within the constrained structural space.

Together, these results show that DropAdd-TS can operate across multiple feature representations while maintaining its diversity-selection behaviour. However, the choice of representation strongly influences the achievable diversity, with sequence identity providing the most informative signal for EC-level functional diversification within homologous protein families.

## Discussion

Selecting representative subsets from large protein sequence datasets remains a central challenge in enzyme discovery when experimental screening capacity is limited. The problem is not simply one of reducing dataset size, but of identifying a fixed-cardinality subset that captures representativeness and diversity to support meaningful downstream experimental screening. Here we show that a bi-level diversity-optimisation framework provides an effective strategy for this task by jointly considering local separation and global dispersion.

The previous MaxSum-based formulation maximises overall dispersion among selected elements but does not prevent the simultaneous inclusion of closely related sequences when they are jointly distant from the remainder of the dataset [1]. This behaviour becomes particularly evident in densely populated sequence spaces containing clusters of highly similar variants. In practice, introducing a MaxMin constraint produced subsets with substantially lower maximum pairwise identities while maintaining comparable or improved recovery of EC annotations. The improved run-to-run stability further suggests that the hierarchical formulation guides the search towards more consistent regions of the solution space than dispersion-based optimisation alone.

A second important observation is that the effectiveness of diversity optimisation depends strongly on how protein similarity is represented. Sequence-based measures produced the strongest separation and the highest recovery of EC-annotated groups, whereas structural similarity compressed the available diversity, likely reflecting the inherent conservation of protein folds within homologous families. Although the optimisation framework itself is representation-independent, the biological interpretation of the selected subsets depends directly on the choice of similarity metric.

From a practical perspective, the ability to select subsets of predefined size makes the approach well suited to plate-based experimental workflows. Unlike clustering-based approaches that rely on similarity thresholds, the optimisation framework directly selects a panel of the desired cardinality while prioritising separation among representatives. This also distinguishes it from coverage-based methods, which prioritise summarising the sequence space and do not guarantee well-separated representatives [19, 12]. In settings where the goal is to sample broadly across a homologous family, this emphasis on separation is particularly useful. The expansion mode further extends this capability by enabling diversity-aware panel growth across successive rounds of screening, supporting iterative metagenomic discovery campaigns that are otherwise often performed as single-pass workflows [17].

Despite these advantages, the optimisation operates entirely on pairwise distances and therefore assumes that the chosen representation reflects meaningful biochemical differences between proteins. The algorithm itself cannot distinguish between biologically competent and compromised candidates if both appear sufficiently distinct in the selected feature space. Consequently, upstream filtering remains important before applying the method to remove truncated sequences, misassembled metagenomic contigs, or variants lacking conserved sequence features critical to the protein family.

A second limitation concerns the functional annotations used to evaluate diversity. EC numbers were used as a proxy for functional diversity because they are widely available and systematically curated. However, EC annotations are relatively coarse descriptors and are not uniformly supported by direct experimental evidence. Because many assignments are inferred from sequence similarity or rule-based annotation pipelines [10], sequence-derived representations may be favoured during benchmarking, partly explaining why sequence identity provided the strongest separation in the present study. In addition, although the method is not restricted to enzymes and can be applied to broader protein families (as illustrated by inclusion of the ABC transporter family), EC numbers primarily reflect enzyme-related functional variation. As a result, the current evaluation remains biased towards enzyme-associated functions.

Future work could extend this framework through more informative similarity representations and richer functional benchmarks. Distance measures that emphasise residues with stronger functional relevance, such as active-site or pocket-proximal regions, may provide more meaningful discrimination than global similarity alone. Hybrid distance matrices that combine multiple representations may further capture complementary aspects of protein variation. Functional benchmarks, such as curated reaction identifiers (e.g. Rhea), experimentally measured activity or binding profiles, or quantitative functional parameters, would provide a stronger basis for evaluating representative selection strategies.

## Conclusion

A bi-level diversity formulation enables the direct selection of diverse, fixed-cardinality subsets while limiting redundancy among closely related sequences. By operating on generalised distance matrices, the DropAdd-TS framework provides a general and context-free strategy for constructing representative protein panels across homologous families, demonstrated here in the context of enzyme-associated functional diversity. The bi-level formulation reduces redundancy among selected sequences by up to 43–46% relative to the previous MaxSum-based approach while maintaining functional coverage, supporting its practical use for constructing diverse screening panels under fixed experimental constraints.

## Abbreviations

MSD: MaxSum Diversity Problem
MMD: MaxMin Diversity Problem
EC: Enzyme Commission
LC: Label Coverage
GSI: Gini-Simpson Index

## Code Availability

The DropAdd-TS implementation and analysis scripts are available at https://github.com/zhenou1/mmdp. A permanent archived version via Zenodo will be released upon publication. Benchmark datasets used in this study are publicly available from the UniProt database.

## Supplementary Information

